# BATTLE-AMP: Benchmarking Antimicrobial Peptide Predictors

**DOI:** 10.64898/2026.06.19.733349

**Authors:** Paulina Szymczak, Adriana Bukała, Wojciech Zarzecki, Michał Sala, Jure Borišek, Setareh Fadavi, Roberto Olayo-Alarcon, Jacek Sroka, Maria Colomé-Tatché, Anna Gambin, Christian L. Müller, Piotr Setny, Ewa Szczurek

## Abstract

As antimicrobial resistance outpaces antibiotic development, antimicrobial peptides (AMPs) have emerged as a promising class of alternative antibacterials, and computational predictors are increasingly used to prioritize AMP candidates. Such predictors are typically evaluated on binary AMP/non-AMP classification, which does not test whether they can identify peptides with clinically relevant potency against specific pathogens. We present BATTLE-AMP, a benchmarking framework that evaluates AMP predictors against experimentally measured minimum inhibitory concentrations (MICs) across clinically relevant bacterial species and strains. We surveyed 48 published methods, finding fewer than 25% reproducible, and benchmarked 10 model families (21 variants) using experimental MIC data, synthetic sequence perturbations, activity cliff analyses, and all-atom molecular dynamics (MD) simulations. Four findings emerge: (i) models trained on MIC data outperform binary classifiers regardless of architecture; (ii) the best model depends on the target pathogen, so model selection must be guided by the biological question; (iii) most models cannot distinguish active peptides from inactive sequences with identical amino acid composition; and (iv) activity cliffs remain unresolved by both machine learning and MD, marking a limit of current computational methods. BATTLE-AMP is released as an open Snakemake framework at https://github.com/szczurek-lab/battleamp-snakemake for benchmarking new models and scoring novel candidate libraries.

---

Antimicrobial resistance (AMR) has outpaced antibiotic development, driving interest in antimicrobial peptides (AMPs) as alternative therapeutics [1]. Modern discovery pipelines, from genomic mining to generative AI [2], can produce millions of candidate sequences *in silico*, but synthesizing and testing them *in vitro* remains slow and expensive. Computational predictors therefore serve as the primary filter, ranking candidates for experimental validation (Figure 1a,b). If these predictors are unreliable, the entire pipeline wastes resources on inactive peptides (Figure 1c).

**Fig. 1.**
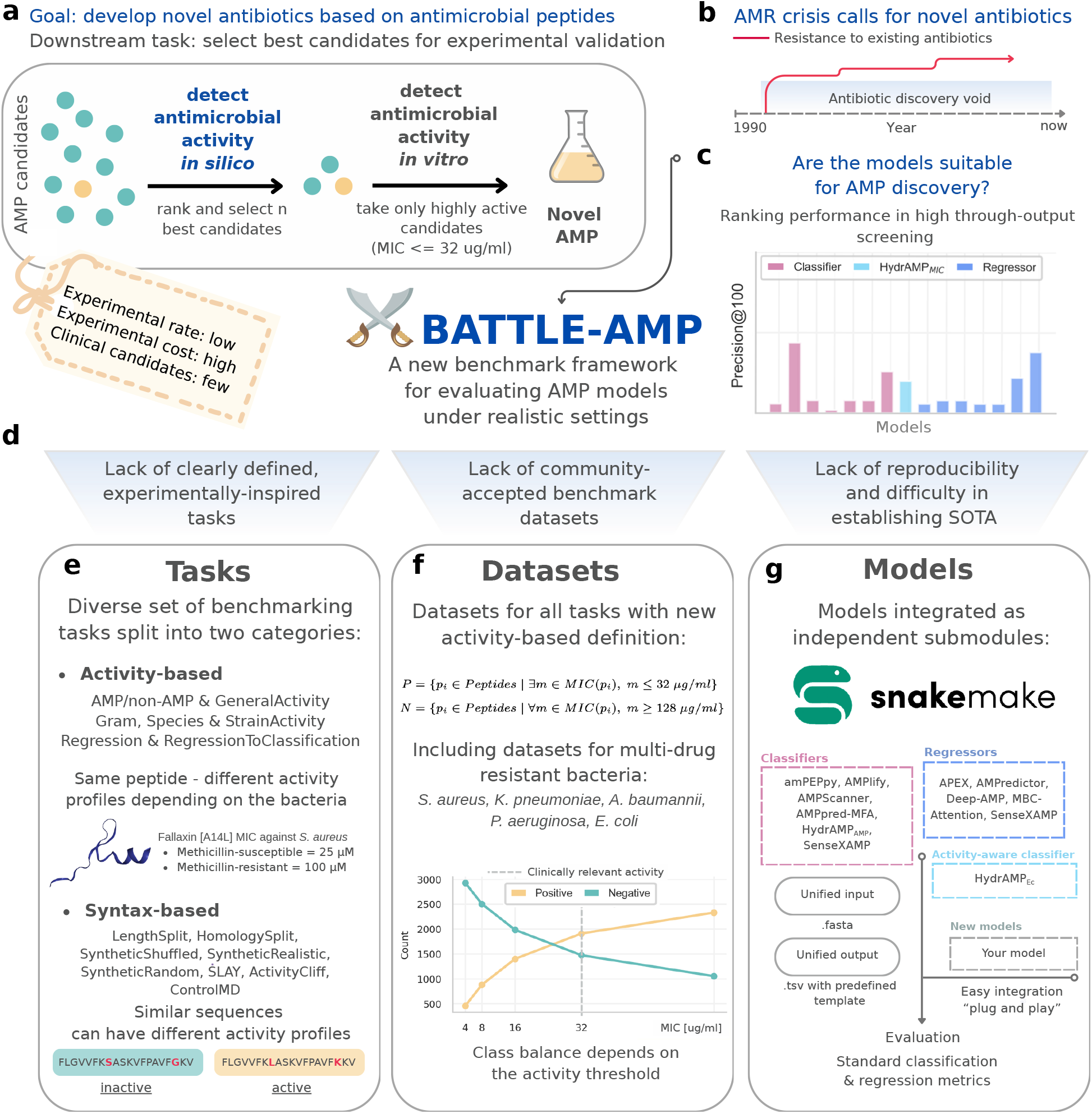
BATTLE-AMP framework overview. (a) AMP discovery pipeline: computational predictors rank candidates *in silico* before experimental validation *in vitro*. (b) The antimicrobial resistance crisis and the antibiotic discovery void. (c) The central question motivating the benchmark: are current models suitable for AMP discovery? Ranking performance in high-throughput screening varies widely across model types. **(d)** Three evaluation gaps motivating the framework: absence of experimentally relevant tasks, lack of accepted benchmark datasets, and poor model reproducibility. (e) The 15 benchmarking tasks, split into activity-based evaluations and syntax-based sequence perturbations. (f) Datasets constructed using clinically relevant activity thresholds (active: MIC≤ 32 µg/ml; inactive: MIC 128≥ µg/ml) for five ESKAPE-associated species, with class balance depending on the selected MIC threshold. (g) Integration of 10 model families (21 variants) as independent Snakemake submodules with a unified FASTA/TSV interface.

Most existing predictors are binary classifiers trained to distinguish known AMPs in databases such as DBAASP [3] and dbAMP [4] from generic non-antimicrobial proteins. This framing is poorly suited to drug discovery: a peptide deposited in database as an AMP may lack therapeutically relevant potency against virulent bacteria. A newer class of models addresses this by predicting measured minimum inhibitory concentrations (MICs) against individual bacterial strains, or by classifying activity at strict clinical thresholds [5–7]. Yet no standardized benchmark evaluates these different modeling approaches against true biological efficacy (Figure 1d). Previous benchmarks have addressed parts of this gap. Xu *et al*. [8] compared predictors at scale but only within the binary paradigm. Sidorczuk *et al*. [9] showed that negative-set sampling strategies bias binary benchmarks, making cross-study comparison unreliable. ESCAPE [10] consolidates over 80,000 peptides from 27 databases into a multilabel classification benchmark covering antibacterial, antifungal, antiviral, and antiparasitic classes, but still evaluates against database-derived labels rather than quantitative potency. PepBenchmark [11] standardizes datasets and evaluation protocols across peptide ML more broadly, though its AMP component relies on a smaller dataset than DBAASP and does not include species-resolved MIC prediction. QMAP [12] introduced homology-aware splits for MIC regression but pools predictions across all bacterial targets into a single score, obscuring whether a model can rank candidates against a specific pathogen.

We present BATTLE-AMP (Figure 1), a benchmarking framework that evaluates AMP predictors against experimentally measured MICs across clinically relevant bacterial species and strains (Figure 1e,f). We surveyed 48 published methods, found fewer than 25% reproducible, and benchmarked 10 model families (21 variants; Figure 1g) using experimental MIC data, synthetic sequence perturbations, activity cliff analyses, and all-atom molecular dynamics (MD) simulations. The framework is implemented as an open Snakemake workflow [13] and is available at Github, enabling practitioners to integrate new models and score novel candidate libraries.

## 2 Results

### 1.1 The AMP/non-AMP task inflates apparent model performance

We first ask whether models that perform well on conventional `AMP/non-AMP` classification also identify peptides with clinically relevant potency. To test this, we compare model performance on the standard `AMP/non-AMP` task with performance on `GeneralActivity`, where labels are based on experimentally measured MIC (active: ≤32 µg/ml; inactive: ≥128 µg/ml).

On the standard `AMP/non-AMP` task, most models achieve strong discrimination (Figure 2a; Table S4).

**Fig. 2.**
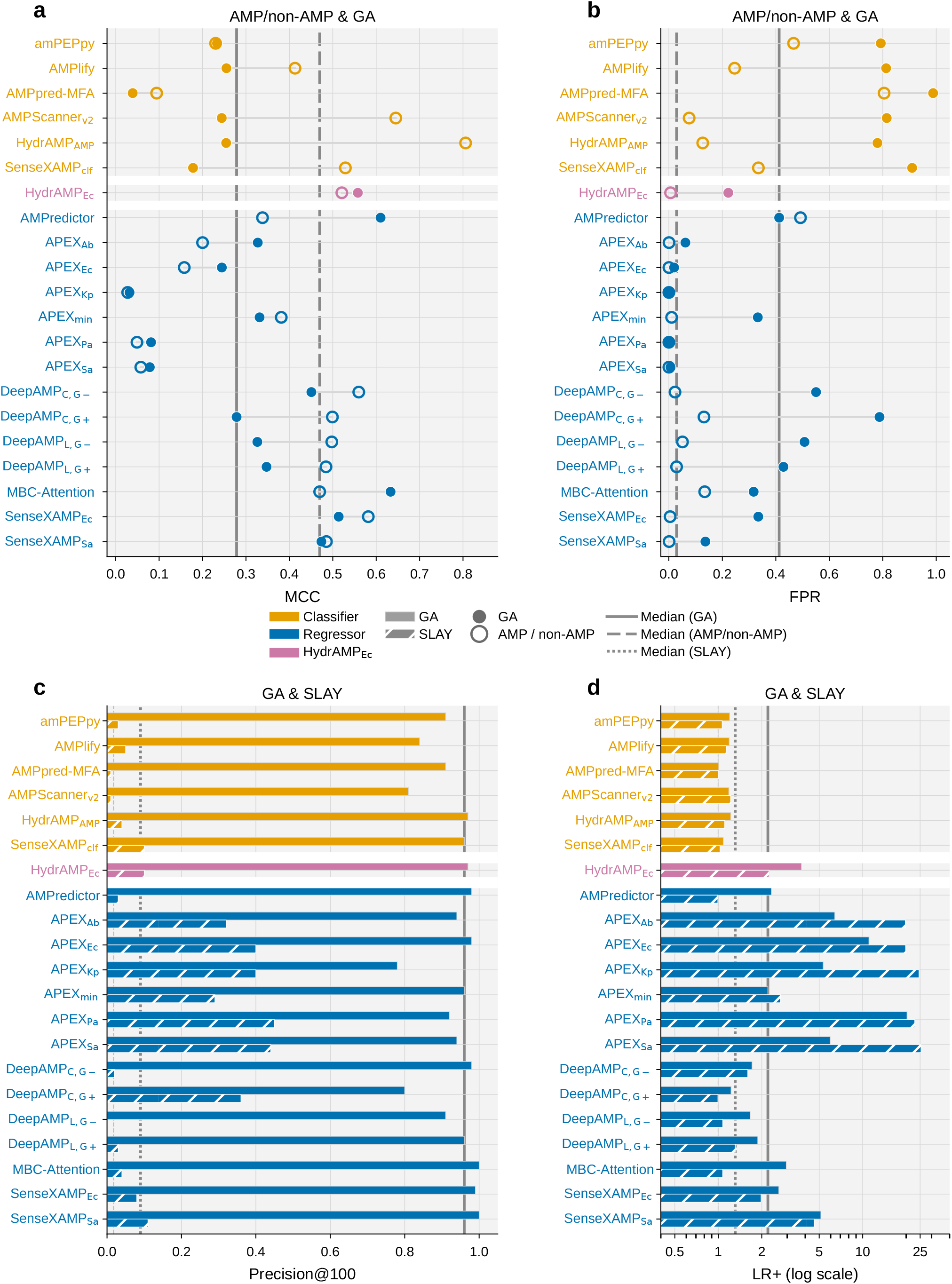
Activity-based benchmarking reveals performance inflation in AMP prediction models. Models are ordered and color-coded by type throughout all panels: classifiers (orange), HydrAMP_Ec_ (pink), and regressors (blue). **(a)** Matthews Correlation Coefficient (MCC) and **(b)** false positive rate (FPR), each shown per model on the `AMP/non-AMP` task (open circle) and on `GeneralActivity` (filled circle); the two markers for each model are connected by a horizontal line to highlight the per-model performance shift between tasks. Vertical reference lines mark the median across models per task: solid for `GeneralActivity`, dashed for `AMP/non-AMP`. **(c)** Precision@100 and **(d)** positive likelihood ratio (LR+, log scale), each shown per model as two bars: a solid bar for `GeneralActivity` and a hatched bar for `SLAY` (*n* ≈ 438,000, ∼ 1.8% actives). Vertical reference lines mark the median across models per task: solid for `GeneralActivity`, dotted for `SLAY`. Regression models were included by binarizing their continuous predictions at clinical thresholds: peptides with MIC≤ 32 µg/ml were labeled active, those with MIC ≥128 µg/ml inactive, and peptides in the intermediate grey zone were excluded.

When evaluation shifts to `GeneralActivity`, which requires experimentally measured potency (MIC≤ 32 µg/ml), typically performance drops sharply (Table S5). The three top-ranked classifiers on `AMP/non-AMP`, HydrAMP_AMP_ (MCC 0.81), AMPScanner_v2_ (0.65), and SenseXAMP_clf_ (0.53), fall to 0.25, 0.24, and 0.18 MCC on `GeneralActivity`, respectively. The only models, for which MCC improves on `GeneralActivity` compared to `AMP/non-AMP` task, are MBC-Attention, AMPPredictor and HydrAMP_Ec_, all trained on quantitative activity data rather than binary AMP/non-AMP labels. False positive rates (FPRs) on `GeneralActivity` exceed 78% for all six classifiers trained on binary AMP/non-AMP labels (Figure 2b). Models trained on measured activity reduce this to 22% (HydrAMP_Ec_) and 14% (SenseXAMP_Sa_); the APEX species-specific regressors achieve near-zero FPRs.

Some evaluation peptides may overlap with model training sets. Of the 4,355 `GeneralActivity` peptides, 773 (18%) appear in AMP databases used to train classifiers, 127 with discordant labels (Supplementary Figure S2). We retain this overlap deliberately, as it penalizes models that call inactive peptides active simply because they appeared in a curated AMP list. To rule out leakage as a driver of the results above, we evaluate all models on the `SLAY` dataset [14]: ∼438,000 random peptides (∼1.8% actives) sharing no sequences with any model’s training data (full per-model metrics in Table S6).

Precision@100 and positive likelihood ratio (LR+) measure how these FPR differences affect a practical screening scenario, on both `GeneralActivity` and the leakage-free `SLAY` set (Figure 2c–d). On `GeneralActivity` (∼78% actives), most activity-trained models reach Precision@100 near 1.0, with LR+ values of 6–20 for the APEX regressors. `SLAY` (∼ 1.8% actives) better approximates the class imbalance of a real screening library. Here, classifiers trained on AMP/non-AMP labels fall to LR+ near 1.0: a positive prediction carries no information about true activity. AMPlify yields 5 actives per 100 top-ranked candidates, and MBC-Attention only 4. A simple physicochemical filter (net charge ≥ +3, hydrophobic moment above the library median) achieves Precision@100 of 0.12 (LR+ 7.5), outperforming every classifier including HydrAMP_Ec_. Only the APEX species-specific regressors exceed this baseline, reaching 40–45 actives per 100 candidates (LR+ 20–25). Overall, models that excel at `AMP/non-AMP` classification do not necessarily identify potent peptides: they produce high FPRs when evaluated against clinically relevant MIC thresholds and offer little to no enrichment over random selection in a screening setting.

### 1.2 Target bacteria shapes prediction difficulty

Most classifiers were trained to distinguish AMPs from non-AMPs, not to predict activity against specific pathogens, or were trained only on a limited set of strains. In practice, however, these models are often the only available filter when selecting candidates for strain-specific experiments, and cross-strain correlations in antimicrobial activity make this a reasonable strategy. Our species- and strain-resolved evaluation measures what signal each model provides for this purpose.

Figure 3a shows MCC for all models across classification tasks of increasing biological specificity. No single model dominates: the best performer changes with the target. Gram− targets are easier than Gram+. On Gram−, four models exceed MCC 0.5, led by MBC-Attention (0.69) and AMPredictor (0.65) (Table S7). On Gram+, only SenseXAMP_Sa_ exceeds 0.5 (0.60), while MBC-Attention drops to 0.42 (Table S8). The two groups have comparable sizes (*n* = 4,012 vs. 3,200), so the gap reflects biological difficulty, not statistical power. Among the APEX regressors, APEX_Ab_ scores highest on four of five species, including *S. aureus* (0.38) and *P. aeruginosa* (0.43). At the strain level, prediction of resistant phenotypes degrades sharply: on susceptible *S. aureus* ATCC 25923 the best model reaches MCC 0.56, but on methicillin-resistant ATCC 33591 and ATCC 43300 the median MCC across models drops to 0.12 and 0.07. SenseXAMP_Sa_ is the exception (MCC 0.58 on both resistant strains), likely because its *S. aureus*-specific training data capture within-species variation that transfers across resistance backgrounds.

**Fig. 3.**
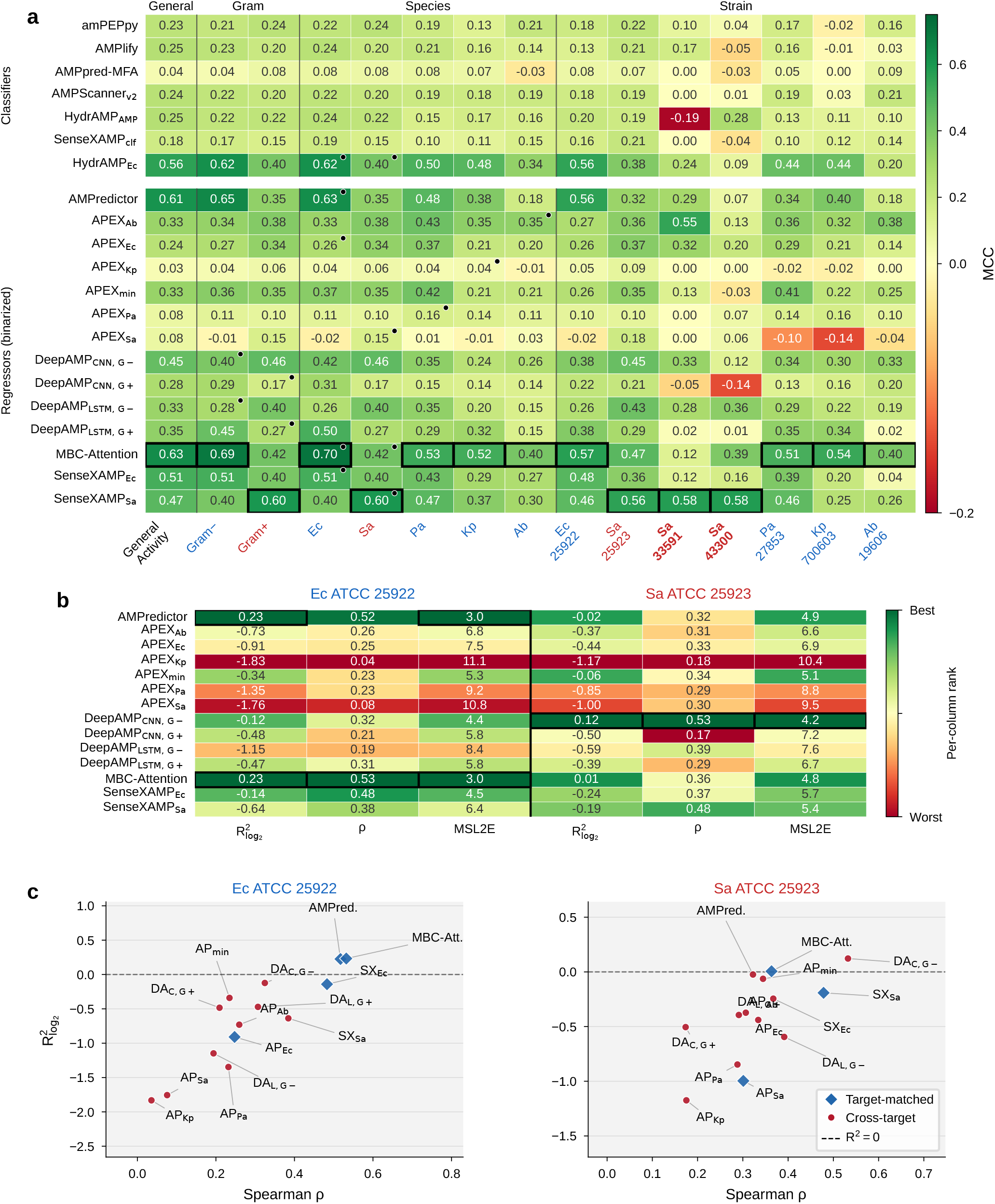
Prediction difficulty varies with target bacteria and favors target-matched training. Throughout the figure, bacterial targets are color-coded by Gram stain (red = Gram-positive, blue = Gram-negative); bold labels indicate multidrug-resistant (MDR) strains. **(a)** MCC heatmap across classification tasks of increasing biological specificity: general AMP detection, Gram type, species, and reference strains. Models are grouped into classifiers (top) and binarized regressors (bottom), separated by a white gap; boxed cells mark the column-wise best; black dots indicate target-matched models. **(b)** Regression performance on *E. coli* ATCC 25922 and *S. aureus* ATCC 25923 (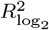, Spearman *ρ*, MSL2E); cells colored by per-column rank (green = best, red = worst). **(c)** 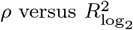 for each regressor on the same two targets; blue diamonds = target-matched models, red circles = cross-target; dashed line marks *R*^2^ = 0. Model abbreviations: AP = APEX, SX = SenseXAMP, DA_L_ = DeepAMP (LSTM), DA_C_ = DeepAMP (CNN), AMPred. = AMPredictor, MBC-Att. = MBC-Attention; subscripts denote bacterial target (Ec = *E. coli*, Sa = *S. aureus*, Pa = *P. aeruginosa*, Kp = *K. pneumoniae*, Ab = *A. baumannii*, min = minimal inhibitory concentration optimized) and, for DeepAMP, Gram type (G+, G*−*).

We note that Precision@*k* values reported in the supplementary tables (Tables S9–S20) are partly inflated by base rate: small species-resolved tasks contain 60–90% active peptides, so even uninformative rankings achieve P@100 above 0.6, and general regressors such as MBC-Attention occasionally outrank target-matched APEX variants on this metric without providing stronger discrimination (cf. MCC and LR+ in the same tables, and the SLAY screening results in Section 1.1, where the realistic ∼1.8% active rate makes P@100 a meaningful discrimination metric).

MIC `Regression` results for *E. coli* ATCC 25922 (*n* = 2,531) and *S. aureus* ATCC 25923 (*n* = 1,568) are shown in Figure 3b (full metrics in Table S21; predicted-versus-observed scatter plots in Supplementary Figures S4 and S5). MBC-Attention and AMPredictor lead on *E. coli* (*ρ* = 0.53 and 0.52; 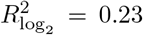 for both). On *S. aureus*, performance drops across all models. Surprisingly, the only model with a clearly positive 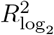_, G*−*_ (0.12), a model trained on Gram−-data. The remaining models fall at or below zero.

SenseXAMP_Sa_, the best-performing model on *S. aureus* in the classification tasks, achieves the second-highest rank correlation here (*ρ* = 0.48) but negative 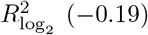, meaning it ranks peptides reasonably but cannot predict their MIC values. APEX models show the same pattern more starkly: they rank well as binarized classifiers yet are among the worst regressors, confirming that their predictions preserve rank order but not quantitative accuracy. The best models err by ~3 log_2_ dilution steps on *E. coli* (MSL2E 3.0) and 4–5 steps on *S. aureus* (MSL2E 4.2–5.4), a roughly 8-to 32-fold error in predicted MIC. This suffices for coarse active/inactive triage but not for dose estimation.

The *ρ* versus 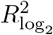 scatter plots (Figure 3c) separate regressors into target-matched (trained on data from the species being evaluated; blue diamonds) and cross-target (trained on a different species; red circles). On both targets, target-matched regressors achieve higher rank correlation and higher variance explained, indicating that training on MIC data from the target species is more informative than training on a different species. However, 11 of 14 regressors still fall below 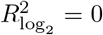, meaning they explain less variance than predicting the evaluation-set mean. Only MBC-Attention and AM Predictor exceed this threshold on *E. coli*, and only Deep AMP _CNN, G*−*_ and marginally MBC-Attention 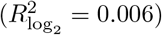 on *S. aureus*. Current models can rank peptides by potency but cannot reliably predict absolute MIC values.

### 1.3 Robustness evaluation reveals reliance on compositional rather than sequence features

Three synthetic negative sets of increasing difficulty (synthetic decoys) test whether models have learned sequence-order features or merely amino acid composition (Figure 4a). `SyntheticRandom` sequences (uniform amino acid sampling) diverge from active peptides in charge and hydrophobic moment, making them trivially separable. `SyntheticRealistic` sequences (matching global AMP amino acid frequencies) close the gap in charge and hydrophobicity but still differ in composition compared to known AMPs. `SyntheticShuffled` sequences (persequence permutations of real actives) preserve exact amino acid composition while destroying residue order, and are therefore compositionally indistinguishable from their parent actives (full physicochemical distributions in Supplementary Figure S3).

**Fig. 4.**
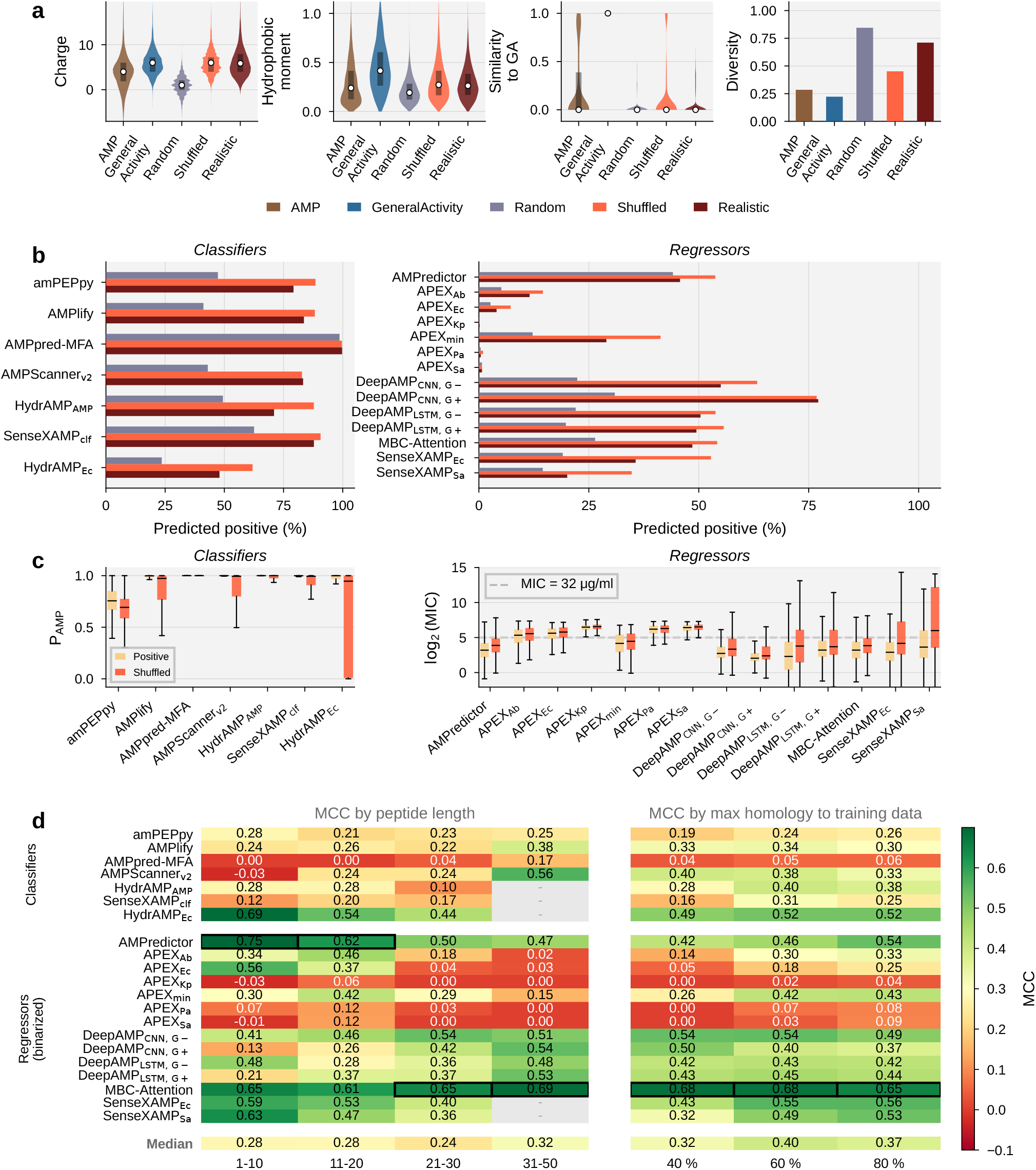
Most models cannot distinguish active peptides from compositionally identical shuffled sequences. **(a)** Physicochemical properties (net charge, hydrophobic moment), sequence similarity to `GeneralActivity`, and sequence diversity for the five evaluation sets: AMP, `GeneralActivity`, `SyntheticRandom`, `SyntheticShuffled`, and `SyntheticRealistic`. **(b)** FPR on `SyntheticRandom`, `SyntheticShuffled`, and `SyntheticRealistic` decoys for classifiers (left) and regressors (right). `GeneralActivity` positives (grey) shown for reference. **(c)** Score distributions for `GeneralActivity` positives versus Shuffled negatives. Left: predicted AMP probability (*P*_AMP_) for classifiers. Right: predicted log_2_(MIC) for regressors; dashed line marks MIC = 32 µg/ml. **(d)** MCC on `GeneralActivity` split by peptide length (left; 1–10, 11–20, 21–30, 31–50 residues) and by maximum sequence homology to training data (right; CD-HIT clustering at 40%, 60%, and 80% identity). Bottom row: median across all models. Black boxes mark the best value per column.

FPR increases sharply as the synthetic negatives become harder to distinguish from real actives (Figure 4b). On `SyntheticRandom` negatives, most models reject the decoys. On the `SyntheticRealistic` set, binary AMP/non-AMP classifiers already fail: AMPlify reaches FPR 0.79, AMPpred-MFA 1.00, and SenseXAMP_clf_ 0.84. On the `SyntheticShuffled` set, where negatives are compositionally identical to their parent actives, nearly all models fail as well. Nine of 21 variants exceed FPR 0.80, including the best-performing models on `GeneralActivity` (MBC-Attention 0.87, AMPredictor 0.89), and no model achieves MCC above 0.17 (per-decoy-set metrics in Tables S22–S24). The score distributions (Figure 4c) show that most classifiers assign nearly identical *P*_AMP_ to actives and their shuffled counterparts, and most regressors predict nearly identical MIC for both. The only exceptions are APEX_Kp_, APEX_Pa_, and APEX_Sa_ (FPR 0.00, 0.01, 0.01), which correctly assign higher MIC to shuffled sequences. Current models thus operate primarily on amino acid composition rather than on residue order or amphipathic structure.

Two additional splits of the `GeneralActivity` task test whether model performance depends on peptide length or on similarity to training data (Figure 4d). Across four length bins (1–10, 11–20, 21–30, 31–50 residues), the median MCC shows no clear trend (0.24–0.32; per-bin metrics in Tables S25–S28), though coverage varies across bins because several models accept only peptides below 25–50 aa. On the shortest peptides (1–10 aa), most models fall below MCC 0.30; exceptions are AMPredictor (0.75), HydrAMP_Ec_ (0.69), MBC-Attention (0.65), and SenseXAMP_Ec_ (0.59). The `HomologySplit` tests a different question: how much performance depends on memorizing training sequences. Restricting evaluation to CD-HIT cluster representatives at 80%, 60%, and 40% identity progressively removes peptides similar to those seen during training. The median MCC drops from 0.37 to 0.32 (per-threshold metrics in Tables S29–S31). Most models degrade, with AMPredictor falling from 0.54 to 0.42 and HydrAMP_AMP_ from 0.38 to 0.28. MBC-Attention is the exception (0.65 at 80%, 0.68 at 40%). Together with the synthetic decoy results, these experiments show two distinct limitations: most models depend on amino acid composition rather than residue order, and their performance degrades as test sequences diverge from training data.

### 1.4 Activity cliff pairs challenge both ML and molecular dynamics simulations

Activity cliff pairs test whether models can detect functional differences between nearly identical sequences. We assembled 50 pairs from DBAASP (median pairwise identity 86%, median edit distance 2 substitutions) and compare them against the `ControlMD` set of 40 diverse peptides (median pairwise identity 17%, median edit distance 18 substitutions). On both sets, we evaluate ML regressors from the preceding sections alongside descriptors from all-atom MD simulations (500 ns per peptide, five 100 ns blocks; Section S6): membrane association (S-score), local bilayer deformation (ZP-scores), backbone helicity (H-scores), and peptide burial depth (BUR-scores).

On the `ControlMD` set both MD descriptors and regressors discriminate active from inactive peptides effectively (Figure 5a–d). S-score distributions separate clearly across all five simulation blocks (Figure 5a), yielding AUC values of 0.80 to 0.85 (Figure 5b). Burial metrics—maximum peptide penetration into the bilayer (PeptMax), maximum water penetration near the peptide (WatMax), and their difference (WatRel)—perform comparably, while among helicity descriptors only H9 (C-terminal helicity; AUC 0.72–0.82) is discriminative. However, this signal is confounded by overrepresentation of C-terminally amidated peptides among actives (Fisher exact test, *p* = 0.044), and stratified analysis confirms that H9 loses significance once amidation is controlled for (Supplementary Section S6). ZP-scores measure how far individual phospholipid phosphate groups are displaced along the membrane normal by the peptide. AUC increases with displacement magnitude for negative ZP (membrane thinning), reaching ∼ 0.85 in block 5 (Figure 5c), but not for positive ZP (thickening), consistent with the thinning mechanism of cationic AMPs. Regressors show even stronger separation, with Spearman *ρ* between predicted log_2_(MIC) and true activity ranging from −0.60 (APEX_min_) to −0.82 (SenseXAMP_Ec_; Figure 5d).

**Fig. 5.**
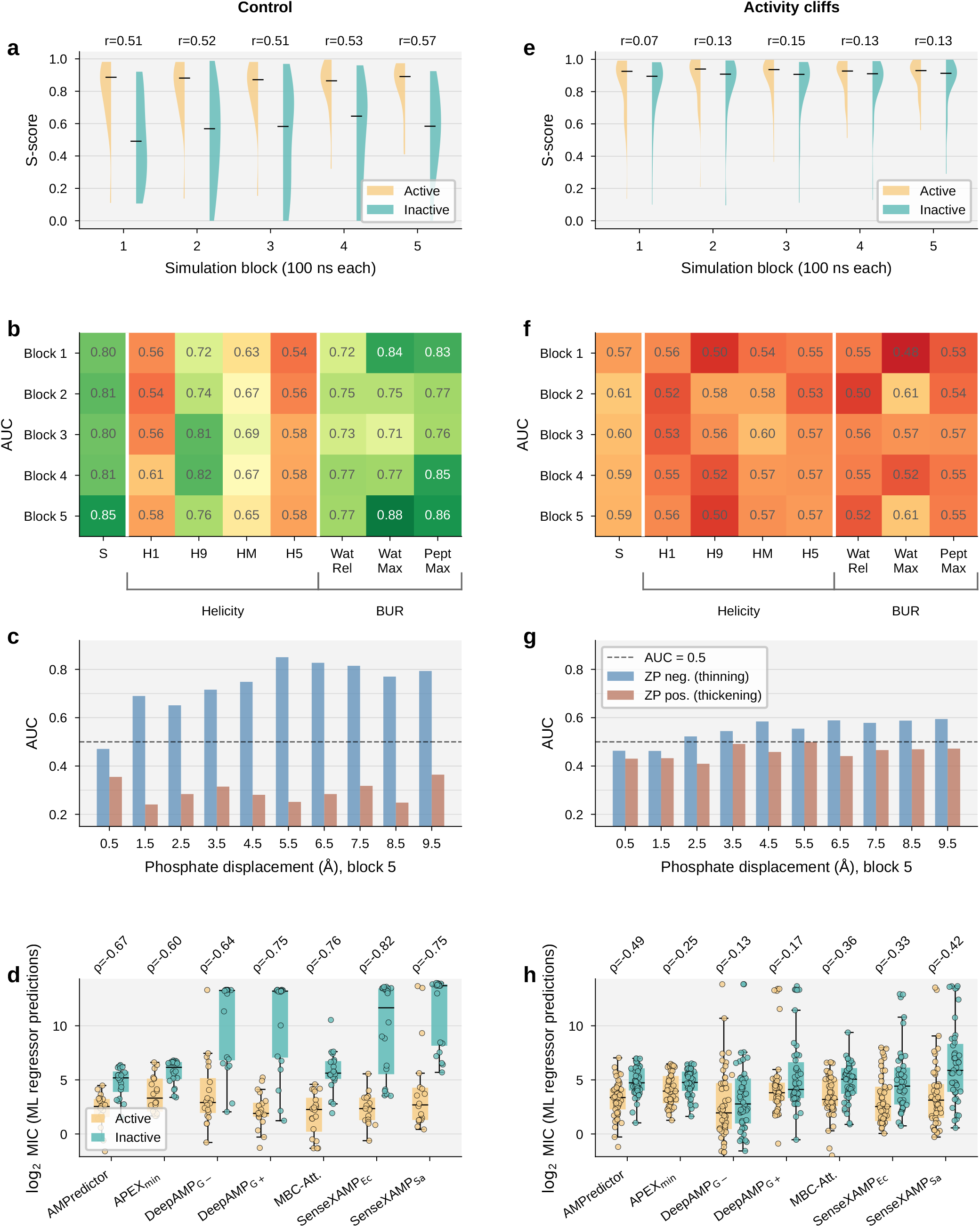
Both MD descriptors and ML regressors discriminate active from inactive peptides on the control set but not on activity cliff pairs. Left column (a–d): `ControlMD` (40 diverse peptides, median pairwise identity 17%). Right column (e– h): `ActivityCliff` set (46 pairs, median pairwise identity 86%). **(a, e)** S-score distributions for active and inactive peptides across five 100 ns simulation blocks; Pearson *r* annotated per block. **(b, f)** AUC heatmaps by MD descriptor family (S-score, helicity H1/H9/HM/H5, burial WatRel/WatMax/PeptMax) and simulation block. Green: AUC *>* 0.70; red: near chance. **(c, g)** ZP-score AUC as a function of phosphate displacement magnitude (block 5). Blue: membrane thinning; orange: membrane thickening. Dashed line: AUC = 0.50. **(d, h)** Predicted log_2_(MIC) from ML regressors for active (orange) and inactive (teal) peptides; Spearman *ρ* annotated per model. Lower MIC = higher potency).

On the `ActivityCliff` set, both approaches fail (Figure 5e–h). S-score distributions for active and inactive partners overlap almost entirely, with Pearson *r* dropping to 0.07–0.15. AUC values across all descriptor families fall to 0.48–0.61, indistinguishable from chance. The relationship between ZP-score AUC and displacement magnitude seen on the control set disappears: AUC remains near 0.50 at all magnitudes (Figure 5g). ML regressors fail in the same way (Figure 5h): predicted MIC distributions for active and inactive cliff partners overlap extensively. That both MD and ML fail on the same pairs suggests that activity cliffs mark a limit of current computational predictability, not a limitation specific to either method.

## 2 Discussion

Via activity-grounded evaluation, we established that training supervision, not model architecture, is the primary determinant of predictive performance: models trained on quantitative MIC data consistently outperform classifiers trained on membership labels, which exhibit false positive rates exceeding 78% and fail to outperform a filter based on charge and hydrophobic moment alone on realistic screening data. This conclusion is consistent with recent findings that simple molecular fingerprint models match or exceed deep learning classifiers across multiple binary AMP benchmarks [15], confirming that architecture alone does not explain performance differences.

Crucially, no single model performs best across all targets, with Gram+ targets proving systematically harder to predict than Gram− . Two factors likely contribute: public databases skew heavily Gram − (*E. coli* alone dominates DBAASP MIC entries), so models see fewer Gram+ training examples; and the thick peptidoglycan cell wall of Gram+ bacteria presents a qualitatively different barrier from the Gram− outer membrane, so sequence-derived features that predict membrane disruption transfer imperfectly across the two groups. Methicillin-resistant strains remain largely intractable, in part due to the scarcity of publicly available MIC measurements against resistant isolates. Beyond varying target difficulty, synthetic decoys experiments reveal that most models have learned amino acid composition statistics rather than sequence-order features, explaining the performance inflation observed under conventional binary evaluation. Finally, activity cliff pairs remain unresolved by both ML regressors and all-atom MD simulations, indicating a fundamental limit of current computational approaches rather than a shortcoming of either paradigm alone.

These conclusions are, however, subject to several constraints. First, BATTLE-AMP evaluates pretrained tools as-is, reflecting realistic practitioner use but preventing full decoupling of architecture from training data. Additionally, activity labels aggregate MIC measurements across laboratories; because MIC values for the same peptide-organism pair can vary by several dilution steps between studies, conservative binarization thresholds reduce but do not eliminate this noise. The MD simulations effectively consider a single peptide molecule on a symmetric bilayer composed of POPE and POPG lipids, a common minimal model of the bacterial inner membrane, over relatively short timescales, thereby omitting cooperative peptide effects on large-scale membrane remodeling, Gram+ cell-wall architecture, and intracellular mechanisms. Furthermore, the benchmark does not yet address toxicity prediction, and covers only the 12 of 48 surveyed tools with accessible, functional code.

Despite these limitations, the results support three concrete recommendations for wet-lab practitioners. First, models trained on species-resolved MIC data should be preferred over AMP/non-AMP classifiers; however, because *E. coli* and *S. aureus* dominate public databases, aggregate performance may not generalize to less-studied pathogens. Second, when screening large libraries where false positives are costly, the APEX species-specific regressors offer 20-to 25-fold enrichment over random selection (Precision@100 of 0.93–0.97 against a background positive rate of 4%). Third, activity cliff pairs, candidates near-identical to known sequences but with differing activity profiles should be flagged for experimental validation, as no current computational method resolves these cases.

The most impactful near-term improvements for future model development are: training on curated, species-resolved MIC data and using composition-preserving shuffled sequences as hard negatives during training, which would directly penalize the compositional shortcuts identified here. Moving beyond compositional dependence will additionally require incorporating structural information through predicted conformations, dynamics-derived features, or structure-aware embeddings. Activity cliffs represent the most pressing unsolved challenge, demanding both richer training data, such as systematic mutagenesis panels with measured MIC, and models capable of learning from minimal sequence variation. More broadly, expanding benchmarks to include toxicity, resistance evolution, and antibiotic synergy would bring evaluation closer to the multi-objective requirements of clinical development. BATTLE-AMP is released as an open Snakemake framework designed to accommodate these extensions.

## 3 Methods

### 3.1 Datasets

All activity datasets were derived from the DBAASP database [3] as (amino acid sequence, MIC value, target bacterial strain) triplets. Sequences were restricted to the 20 standard amino acids (aa) and allowed terminal modifications. Activity measurements were filtered to retain only entries with colony-forming unit (CFU) densities of 10^5^–10^6^ CFU/ml, assay media conforming to standard susceptibility testing protocols (MHB, CAMHB, and related agar/broth formulations), and a measure type of MIC. To construct binary classification datasets, peptides were partitioned into positive (*P*, active) and negative (*N*, inactive) sets based on their minimum MIC across all tested strains:

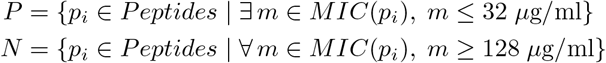

The 32 µg/ml lower bound is a conventional threshold in the AMP literature [5, 16] for clinically relevant potency, and the 128 µg/ml upper bound marks a concentration of negligible therapeutic value; the four-fold gap between them spans two twofold-dilution steps, exceeding typical inter-laboratory MIC variability and ensuring class assignments are robust to single-dilution measurement error. Peptides with intermediate MIC values (MIC ∈ (32, 128) µg/ml, hereafter the grey zone) were excluded to reduce label noise (Figure S1; full rationale and threshold tuning in Supplementary Methods S1). Because conventional classifiers treat any peptide with documented antimicrobial properties as a positive instance regardless of potency, their positive training sets inevitably intersect with our inactive set *N* (Supplementary Figure S2). This choice is deliberate: our framework is built to penalize models that predict weak peptides (MIC ≥128 µg/ml) as active merely because they share generic AMP features. We further define datasets at specific taxonomic levels (Gram type, species, or strain) by filtering peptide subsets accordingly. For example, positives for Gram+ bacteria are peptides active against at least one Gram+ strain.

The `AMP/non-AMP` dataset combines deduplicated positives from dbAMP [4], AMPScanner [17], and DRAMP [18] with non-antimicrobial cytoplasmic SwissProt [19] negatives (8–200 aa); construction protocol in Supplementary Methods S1.

### 3.2 Benchmarking tasks

We constructed 15 benchmarking tasks in two categories (Table 1; dataset sizes in Supplementary Table S1): *activity-based tasks*, which evaluate predictions of antimicrobial potency at increasing taxonomic resolution, and *syntax-based tasks*, which probe model robustness to sequence-level perturbations. `GeneralActivity` evaluates binary activity against any strain; `GramActivity`, `SpeciesActivity`, and `StrainActivity` progressively resolve predictions to Gram type, individual species, and reference strains. `Regression` tasks evaluate quantitative MIC predictions, and `RegressionToClassification` binarizes regressor outputs at the standard thresholds for direct comparison with classifiers.

**Table 1.**
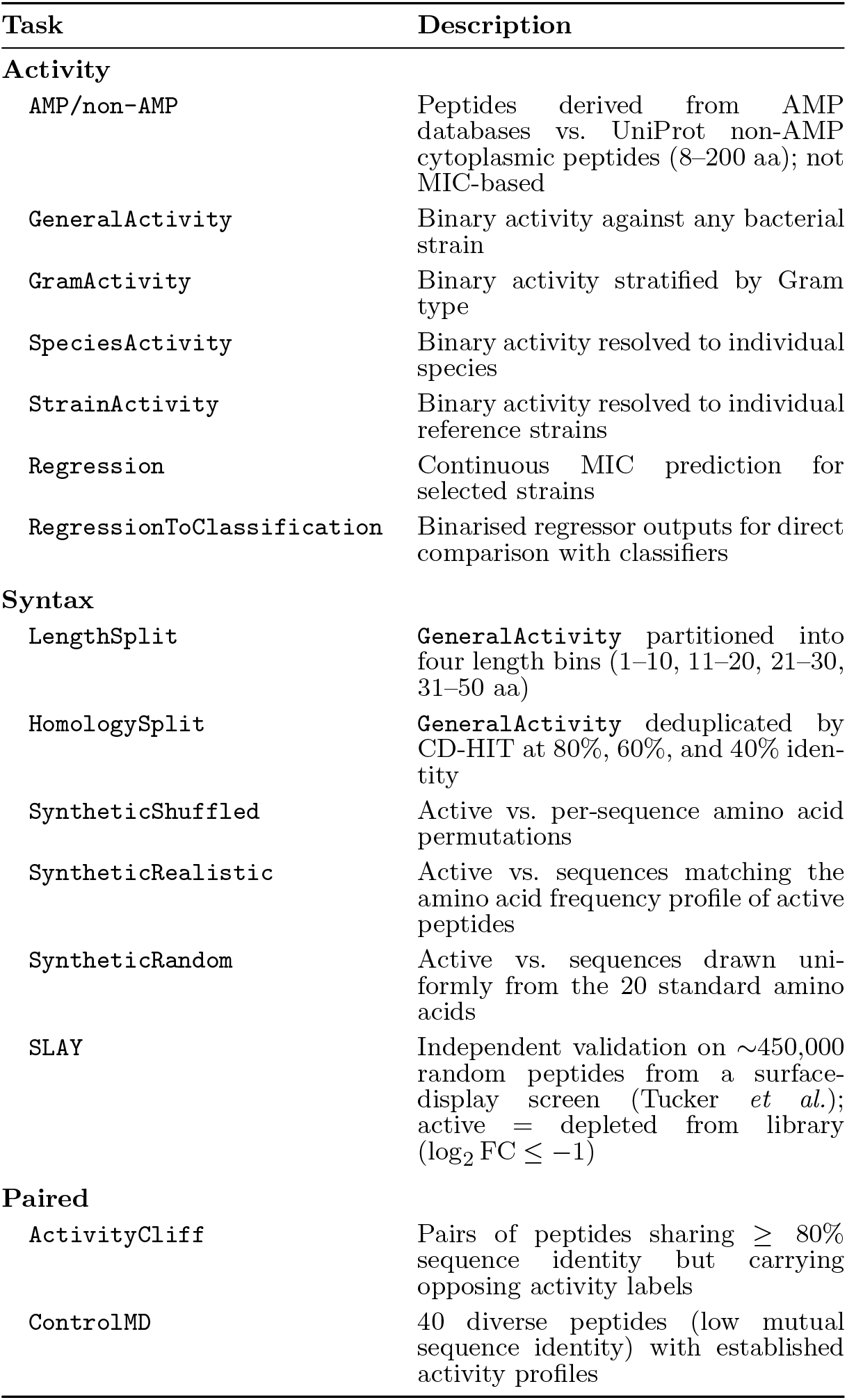
Benchmarking tasks in BATTLE-AMP. Unless otherwise noted, peptides are labeled active (MIC *≤* 32 µg/ml) or inactive (MIC *≥* 128 µg/ml) based on experimentally measured potency.

| Task | Description |
| --- | --- |
| **Activity** |  |
| AMP/non-AMP | Peptides derived from AMP databases vs. UniProt non-AMP cytoplasmic peptides (8-200 aa); not MIC-based |
| GeneralActivity | Binary activity against any bacterial strain |
| GramActivity | Binary activity stratified by Gram type |
| SpeciesActivity | Binary activity resolved to individual species |
| StrainActivity | Binary activity resolved to individual reference strains |
| Regression | Continuous MIC prediction for selected strains |
| RegressionToClassification | Binarised regressor outputs for direct comparison with classifiers |
| **Syntax** |  |
| LengthSplit | **GeneralActivity** partitioned into four length bins (1-10, 11-20, 21-30, 31-50 aa) |
| HomologySplit | **GeneralActivity** deduplicated by CD-HIT at 80%, 60%, and 40% identity |
| SyntheticShuffled | Active vs. per-sequence amino acid permutations |
| SyntheticRealistic | Active vs. sequences matching the amino acid frequency profile of active peptides |
| SyntheticRandom | Active vs. sequences drawn uniformly from the 20 standard amino acids |
| SLAY | Independent validation on $\sim 450,000$ random peptides from a surface-display screen (Tucker *et al.*); active = depleted from library ($\log_2 \text{FC} \leq -1$) |
| **Paired** |  |
| ActivityCliff | Pairs of peptides sharing $\geq 80\%$ sequence identity but carrying opposing activity labels |
| ControlMD | 40 diverse peptides (low mutual sequence identity) with established activity profiles |

Syntax-based tasks evaluate model sensitivity to sequence syntax and compositional biases. `LengthSplit` and `HomologySplit` partition the `GeneralActivity` set by peptide length and by CD-HIT clustering identity (80%, 60%, 40%), respectively, quantifying reliance on training-set length distributions and sequence memorization. Three *Synthetic* subtasks pair active peptides against progressively harder negative sets (hereafter synthetic decoys): `SyntheticRandom` (drawn uniformly over the 20 canonical aa), `SyntheticRealistic` (matching empirical AMP aa frequencies), and `SyntheticShuffled` (randomly permuting each real AMP to preserve exact composition while destroying residue order); physicochemical properties of all three synthetic sets are compared against active and inactive peptides in Supplementary Figure S3. The `SLAY` task provides external validation on ∼450,000 random peptides from a surface-display antimicrobial screen by Tucker *et al*. [14], where peptides that kill the host cell are depleted from the library.

To probe the predictive boundary of current models, we introduce two paired datasets. `ActivityCliff` isolates peptides sharing ≥ 80% sequence identity but carrying opposing activity labels (median pairwise identity 86%, median edit distance 2 substitutions), testing sensitivity to minimal, functionally disruptive mutations. A complementary `ControlMD` set of 40 diverse peptides (median pairwise identity 17%, median edit distance 18 substitutions) with established activity profiles serves as a baseline for MD descriptors, ensuring any loss of performance on activity cliffs reflects genuine task difficulty.

### 3.3 Reproducibility survey and model selection

We surveyed 48 AMP prediction methods published up to March 2024 (Table S2), for each attempting to obtain source code, resolve the computational environment, and run inference on BATTLE-AMP sequences. Of the 48 tools, 25 had code nominally available but only 12 could be executed successfully; two of these (MolE [20] and sAMP-VGG16 [21]) were subsequently excluded for producing constant predictions across all inputs. Dominant failure modes included broken repository links, undocumented preprocessing, and unresolvable dependency conflicts. Adaptation details and the complete survey are provided in Supplementary Methods S2 and Table S2.

From the reproducible subset, we benchmarked 10 base tools yielding 21 model variants. These span random forests trained on physicochemical descriptors (amPEPpy [22]), convolutional and recurrent sequence models (AMP Scanner v2 [17], AMPlify [23], AMPpred-MFA [24], DeepAMP [25], MBC-Attention [6], HydrAMP [16]), graph convolutional networks over protein language (pLMs) model embeddings (AMPredictor [26]), attention-based recurrent ensembles (APEX [5]), and a multi-modal architecture fusing ESM embeddings with structural descriptors (SenseXAMP [27]). Variant definitions and integration details are provided in Supplementary Methods S3.

### 3.4 Evaluation

BATTLE-AMP is a pure inference benchmark: all models are evaluated using their original pretrained weights with no retraining, reflecting the practitioner scenario of applying a published tool to a novel candidate library. For classification, we report the Matthews Correlation Coefficient (MCC) as the primary metric, alongside FPR, AUROC, AUPRC, partial AUROC at FPR ≤ 0.1 and 0.01, Precision@*k* (*k* = min(100, *n*)), and positive likelihood ratio (LR+). MCC was selected because it accounts for all four confusion matrix entries and remains robust under class imbalance. We additionally report AUPRC and Precision@*k* to cover the complementary screening scenario in which only the top-ranked candidates proceed to validation; these ranking-based metrics are the relevant ones for high-throughput libraries where the absolute threshold matters less than the ordering of candidates.

For `Regression`, we report *R*^2^ on log_2_-transformed MIC, Spearman rank correlation, and RMSL^2^E, with predictions clamped to 0.25–512 µg/ml. Spearman rank correlation was used for MIC predictions, which are ordinal by nature (twofold dilution steps); Pearson correlation was used for continuous MD descriptors. Kendall’s *τ* is additionally reported in the supplementary tables as a tie-robust alternative given the discrete nature of MIC dilution series. Regressors evaluated on classification tasks had their MIC predictions binarized at the standard thresholds (≤32 active, ≥128 inactive), with grey-zone values excluded. Complete metric definitions are given in Supplementary Methods S4.

### 3.5 Framework design

BATTLE-AMP is implemented as a modular, open-source Snakemake workflow [13]. Each model is encapsulated as an independent submodule with a dedicated Conda environment and standardized inference script (FASTA in, TSV out), decoupling model-specific dependencies from the benchmark infrastructure. Practitioners can customize execution via the configuration file to run specific subsets of models, tasks, and metrics. New models can be integrated by providing an environment definition and inference script. Inference time and memory requirements for each tool are summarised in Supplementary Table S3. Architectural details are described in Supplementary Methods S5.

### 3.6 Molecular dynamics simulations

All-atom MD simulations of peptide-membrane interactions were performed on two sets: (i) 50 activity cliff pairs, of which 46 yielded stable trajectories, and (ii) the 40-peptide `ControlMD` set, of which 36 completed successfully. Initial peptide structures were modelled as ideal *α*-helices and placed on opposite sides of a model bacterial membrane (POPE:POPG, 2:1; 180 lipids), yielding two independent peptide trajectories per simulation. Systems were solvated in TIP3P water with 150 mM KCl and simulated at the temperature of 310 K for 500 ns using GROMACS [28] with the CHARMM36m force field [29]. Terminal modifications were applied as recorded in the source databases.

Peptide-membrane interactions were characterized by four descriptor families computed in 100 ns windows: S-scores (membrane contact fraction) [16], ZP-scores (z-axis lipid phosphate displacement capturing local membrane deformation), H-scores (per-residue *α*-helicity via Define Secondary Structure of Proteins (DSSP) [30]), and BUR-scores (peptide and proximal water burial depths). Full simulation protocols and descriptor definitions are provided in Supplementary Methods S6.

## Acknowledgments

This project has received funding from the European Research Council (ERC) under the European Funding Union’s Horizon 2020 research and innovation programme (grant agreement No 810115 – DOG-AMP). JB acknowledges the Slovenian Research and Innovation Agency (ARIS) for financial support (Grant: P1-0017). PSe was supported by the Polish National Science Centre Grant 2020/38/E/ST4/00319. This research was carried out with the support of the Interdisciplinary Centre for Mathematical and Computational Modelling University of Warsaw (ICM UW) under computational allocation no G100-2273. We gratefully acknowledge Poland’s high-performance Infrastructure PLGrid ACK Cyfronet AGH, for providing computer facilities and support within computational grant no PLG/2024/017428. R.O.A was funded by a grant awarded to C.L.M. for the StressRegNet consortium within the Bavarian research network bayresq.net funded through the Bavarian State Ministry of Science and Arts, Germany.

## Author contributions statement

CRediT: PSz conceptualization, data curation, methodology, software, visualization, writing – original draft, review & editing; AB conceptualization, methodology, software, visualization, writing - review & editing; WZ methodology, software, writing - review & editing; MS investigation; JB investigation; SF investigation; ROA software; JS software; MCT supervision; AG supervision; CLM supervision; PSe methodology, investigation, supervision; ES conceptualization, methodology, supervision, funding acquisition, writing - review & editing.

## Competing interests

Szczurek lab receives funding from Merck Healthcare KGaA. The other authors declare no competing interests.

## Ethics approval

This study did not involve human participants, identifiable human data or tissue, or animal experiments. All analyses were performed on previously published, publicly available peptide sequence and antimicrobial activity data. Ethics approval was therefore not required.

## Data availability

All benchmark datasets, evaluation code, model integration scripts, and the complete Snakemake workflow are available at https://github.com/szczurek-lab/battleamp-snakemake. Raw activity data were obtained from the DBAASP database [3]; the `SLAY` dataset from GEO accession GSE94529.

## Notes

https://github.com/szczurek-lab/battleamp-snakemake

### Competing Interest Statement

The authors have declared no competing interest.

## References

[1] Hancock, R. E. & Sahl, H.-G. Antimicrobial and host-defense peptides as new anti-infective therapeutic strategies. Nature Biotechnology 24, 1551–1557 (2006).

[2] Szymczak, P. et al. AI-driven antimicrobial peptide discovery: mining and generation. Accounts of Chemical Research 58, 1831–1846 (2025).

[3] Pirtskhalava, M. et al. DBAASP v3: database of antimicrobial/cytotoxic activity and structure of peptides as a resource for development of new therapeutics. Nucleic Acids Research 49, D288–D297 (2021).

[4] Jhong, J.-H. et al. dbAMP 2.0: updated resource for antimicrobial peptides with an enhanced scanning method for genomic and proteomic data. Nucleic Acids Research 50, D460–D470 (2022).

[5] Wan, F., Torres, M. D., Peng, J. & de la Fuente-Nunez, C. Deep-learning-enabled antibiotic discovery through molecular de-extinction. Nature Biomedical Engineering 8, 854–871 (2024).

[6] Yan, J., Zhang, B., Zhou, M., Campbell-Valois, F.-X. & Siu, S. W. A deep learning method for predicting the minimum inhibitory concentration of antimicrobial peptides against Escherichia coli using Multi-Branch-CNN and Attention. mSystems 8, e00345–23 (2023).

[7] Sharma, R. et al. Artificial intelligence-based model for predicting the minimum inhibitory concentration of antibacterial peptides against ESKAPEE pathogens. IEEE Journal of Biomedical and Health Informatics 28, 1949–1958 (2023).

[8] Xu, J. et al. Comprehensive assessment of machine learning-based methods for predicting antimicrobial peptides. Briefings in Bioinformatics 22, bbab083 (2021).

[9] Sidorczuk, K. et al. Benchmarks in antimicrobial peptide prediction are biased due to the selection of negative data. Briefings in Bioinformatics 23, bbac343 (2022).

[10] Ojeda, S. et al. A standardized benchmark for multilabel antimicrobial peptide classification. arXiv preprint arXiv:2511.04814 (2025).

[11] Zhang, J. et al. Pepbenchmark: A standardized benchmark for peptide machine learning. arXiv preprint arXiv:2604.10531 (2026).

[12] Lavertu, A., Corbeil, J. & Germain, P. QMAP: A benchmark for standardized evaluation of antimicrobial peptide MIC and hemolytic activity regression. bioRxiv 2026–02 (2026).

[13] Mölder, F. et al. Sustainable data analysis with Snakemake. F1000Research 10, 33 (2025).

[14] Tucker, A. T. et al. Discovery of next-generation antimicrobials through bacterial self-screening of surface-displayed peptide libraries. Cell 172, 618–628 (2018).

[15] Adamczyk, J., Ludynia, P. & Czech, W. Molecular fingerprints are strong models for peptide function prediction. Bioinformatics btag179 (2026).

[16] Szymczak, P. et al. Discovering highly potent antimicrobial peptides with deep generative model HydrAMP. Nature Communications 14, 1453 (2023).

[17] Veltri, D., Kamath, U. & Shehu, A. Deep learning improves antimicrobial peptide recognition. Bioinformatics 34, 2740–2747 (2018).

[18] Ma, T. et al. DRAMP 4.0: an open-access data repository dedicated to the clinical translation of antimicrobial peptides. Nucleic Acids Research 53, D403–D410 (2025).

[19] Bateman, A. et al. UniProt: the universal protein knowledgebase in 2025. Nucleic Acids Research 53 (2025).

[20] Olayo-Alarcon, R. et al. Pre-trained molecular representations enable antimicrobial discovery. Nature Communications 16, 3420 (2025).

[21] Pandey, P. & Srivastava, A. sAMP-VGG16: Drude polarizable force-field assisted image-based deep neural network prediction model for short antimicrobial peptides. bioRxiv 2023–06 (2023).

[22] Lawrence, T. J. et al. amPEPpy 1.0: a portable and accurate antimicrobial peptide prediction tool. Bioinformatics 37, 2058–2060 (2021).

[23] Li, C. et al. AMPlify: attentive deep learning model for discovery of novel antimicrobial peptides effective against who priority pathogens. BMC Genomics 23, 1–15 (2022).

[24] Li, C., Zou, Q., Jia, C. & Zheng, J. AMPpred-MFA: an interpretable antimicrobial peptide predictor with a stacking architecture, multiple features, and multihead attention. Journal of Chemical Information and Modeling 64, 2393–2404 (2023).

[25] Pandi, A. et al. Cell-free biosynthesis combined with deep learning accelerates de novo-development of antimicrobial peptides. Nature Communications 14, 7197 (2023).

[26] Dong, R. et al. Exploring the repository of de novo-designed bifunctional antimicrobial peptides through deep learning. eLife 13, RP97330 (2025).

[27] Zhang, W., Xu, Y., Wang, A., Chen, G. & Zhao, J. Fuse feeds as one: cross-modal framework for general identification of AMPs. Briefings in Bioinformatics 24 (2023).

[28] Abraham, M. J. et al. GROMACS: High performance molecular simulations through multi-level parallelism from laptops to supercomputers. SoftwareX 1, 19–25 (2015).

[29] Huang, J. & MacKerell Jr, A. D. CHARMM36 all-atom additive protein force field: Validation based on comparison to NMR data. Journal of Computational Chemistry 34, 2135–2145 (2013).

[30] Kabsch, W. & Sander, C. Dictionary of protein secondary structure: pattern recognition of hydrogen-bonded and geometrical features. Biopolymers: Original Research on Biomolecules 22, 2577–2637 (1983).

